# Immune cell *de novo* steroidogenesis regulates inflammation resolution and recovery in acute lung injury

**DOI:** 10.1101/2025.08.29.673039

**Authors:** Hosni A.M. Hussein, Sanu Korumadathil Shaji, Clara Veiga-Villauriz, Soura Chakraborty, Jhuma Pramanik, Fatma Abo Zakaib Ali, Jing Yuan, Esam Khanfar, Ntombizodwa Makuyana, Youssif M. Ali, Qiuchen Zhao, Daniel Hebenstreit, Bidesh Mahata

## Abstract

Effective resolution of inflammation following acute lung infection or injury is critical for restoring immune and tissue homeostasis to ensure functional recovery. Prolonged or unresolved inflammation can impair lung repair, promote fibrosis, and contribute to pulmonary dysfunction. While systemic steroid signalling is known to modulate general immune responses, the specific role of immune cell-mediated steroidogenesis in regulating lung inflammation and repair remains unknown. Here, we show that immune cell *de novo* steroidogenesis is essential for resolving inflammation and promoting recovery in a murine model of acute lung injury. During the resolution phase, steroid-synthesizing immune cells, predominantly basophils, are enriched in the lung. Mice with immune cell-specific ablation of *de novo* steroidogenesis exhibit exacerbated lung injury, impaired resolution of inflammation, and defective tissue repair. These findings reveal a previously unrecognized immunoregulatory function of immune cell-derived steroids and identify immune cell steroidogenesis as a potential therapeutic target for promoting resolution and recovery in inflammatory lung diseases.

## Main

Inflammatory response following lung infection and injury is critical to remove pathogens and dead cells, however persistent inflammation leads to tissue damage and exacerbates the disease outcome ^1, 2, 3, 4^. Unresolved lung inflammation impairs epithelial regeneration, and disrupts the restoration of normal lung architecture, resulting in conditions such as fibrosis or chronic inflammation. Respiratory infections (RIs) caused by viruses and bacteria are the major etiological factors underlying acute lung injury (ALI), posing substantial health and economic hurdles, particularly for critically ill patients ^5^. Severe RIs result in prolonged inflammation and significant loss of the respiratory epithelial cells leading to life-threatening forms of lung failure^2, 6^. Active inflammation resolution and regenerative responses in the lung at the resolution phase following RI are crucial to resolve inflammation, repair damaged tissue, and restore lung homeostasis and function. Individuals who survive RI develop chronic pulmonary diseases if the lung fails to resolve inflammation and regenerate properly ^7, 8, 9, 10, 11, 12, 13^. Immune cells associated with recovery are enriched in the lung during the resolution phase of RI and play a pivotal role in restoring pulmonary homeostasis ^14, 15, 16^. Immune response during the resolution phase of RI stands as the primary factor influencing successful recovery and lung repair ^17^. However, the precise mechanism is unclear.

Steroid hormones are known to regulate immune cell function and inflammation^18^. Interestingly, recent studies have shown that immune cells express the rate-limiting steroidogenic enzyme Cyp11a1 (also known as cytochrome P450 side-chain cleavage enzyme) and are able to synthesize steroids *de novo* ^19, 20, 21^. The role of immune cell steroidgenesis has been characterise in T cells, macrophages and mast cells in regulating immune cell function, particularly in cancer ^22, 23, 24, 25^. However, its role in facilitating resolution of inflammation and recovery after ALI remains unexplored. Thus, in this study we aimed to investigate how this pathway contributes to the restoration of tissue homeostasis and recovery in a model of bacterial lipopolysaccharide (LPS)-induced ALI.

To determine the dynamic of *de novo* steroidogenic immune cells following lung injury, we used *Cyp11a1^mCherry^* reporter mice^19, 23^ in an LPS-induced ALI model (Fig. 1a). Days 3 and 7 were selected as key time points to assess the inflammatory and resolution phases, respectively ^26, 27^. LPS-treated mice exhibited a significant loss of body weight compared to PBS-treated controls (Fig. 1b). LPS treatment typically induces pathological changes in the lung ^28, 29, 30^. To evaluate these changes in our ALI model, tissue histology was examined in PBS-treated group (Fig. 1c), as well as on day 3 (Fig. 1d) and 7 (Fig. 1e) following LPS treatment. H&E staining indicated that lung tissues from LPS-treated mice exhibited significantly higher scores of pathological alterations compared to PBS control group (Fig. 1f-i). Bronchial alteration (Fig. 1f), inflammation (Fig. 1g), inter-alveolar septal thickness (Fig. 1h), and total lesion score (Fig. 1i) were significantly higher on day 3 post LPS-treated mice compared to PBS control group. On day 7 of post-treatment, the injury score appeared significantly reduced when compared to day 3 (Fig. 1c-i). Following LPS treatment, inflammatory immune cells (mainly myeloid cells) accumulate in the lung during the inflammation phase and subsequently diminish during the recovery phase ^31, 32, 33^. We found that the number of neutrophils and interstitial macrophages (IMs) significantly increased in the lung during the inflammatory phase and resolved by day 7 post LPS treatment (Fig. 1j). In contrast, we observed a significant reduction in the number of alveolar macrophages (AMs) on day 3 post-treatment, followed by a gradual recovery during the resolution phase (Fig. 1j). The depletion of AMs correlates with the severity of lung damage, while their replenishment typically occurs during the resolution phase ^34, 35, 36, 37, 38^. Moreover, we observed a marked increase in the number of B cells in the lung on day 3, which continued to rise by day 7 following LPS treatment (Fig. 1j).

**Fig. 1.**
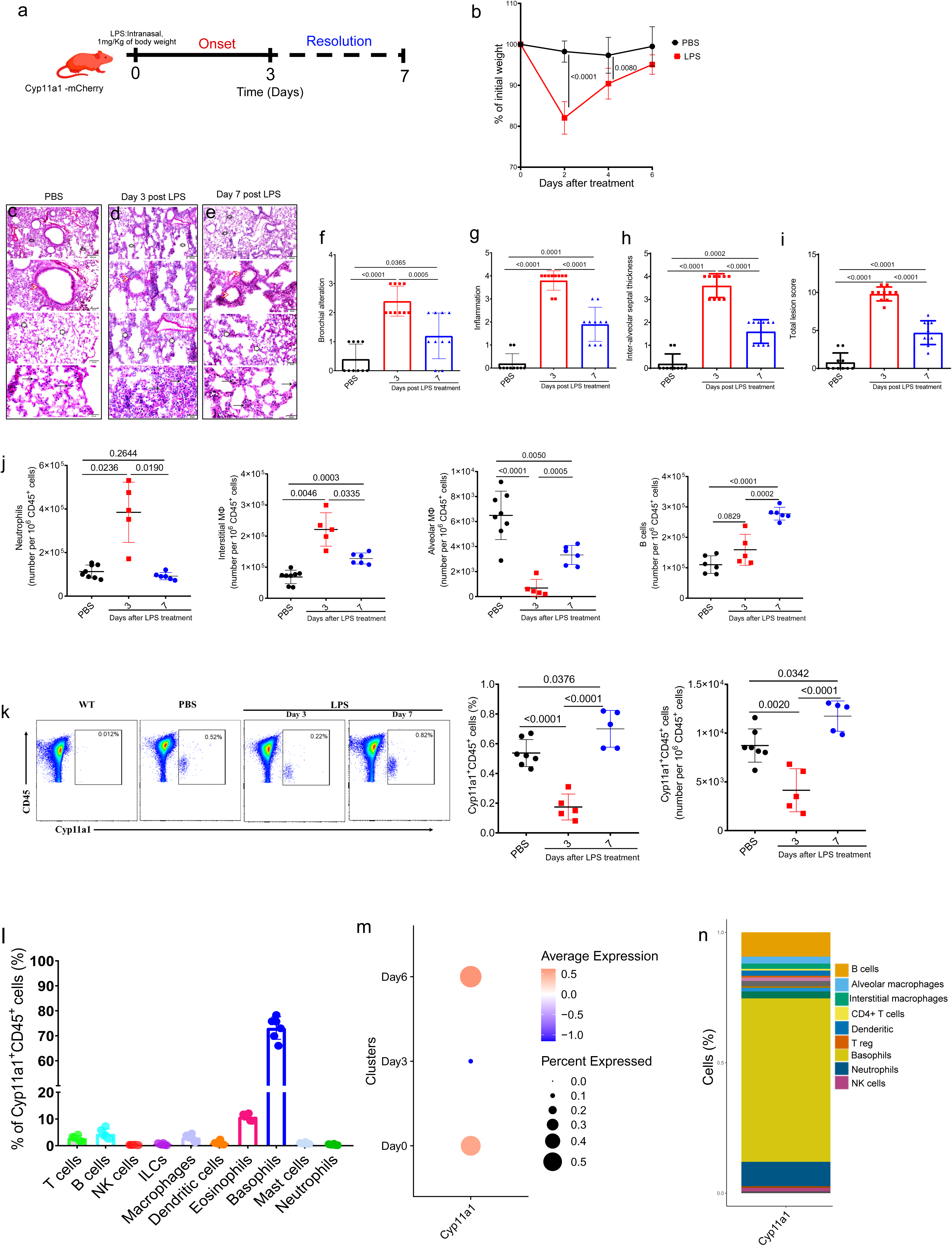
Cyp11a1 expression in immune cells following LPS treatment. **a,** Schematic representation of LPS-induced acute lung inflammation (ALI) experiment in Cyp11a1-mCherry reporter. Cyp11a1^mCherry^ reporter mice were anesthetized and intranasally administered 50 µL of either sterile PBS or LPS. **b**, Body weight changes in LPS or PBS treated mice. (n=5-8) **c-e**, Representative images of hematoxylin and eosin (H&E) staining of lung sections from PBS (n=8) or LPS treated mice on day 3 (n=5) and 7 (n=6) post LPS treatment. **c,** PBS control group: Normal inflated alveoli (stars), normal bronchial epithelium (red arrowheads), normal inter-alveolar septa (white arrowheads), and normal infiltrated alveolar macrophages (normal arrows). **d,** LPS, Day 3 group: alveolar emphysema (stars), moderate peri-bronchial inflammatory cellular infiltration (red arrowheads), marked thick inter-alveolar septa (white arrowheads), and proliferated alveolar epithelial cells mixed with intense infiltrated inflammatory cells (thin arrows). **e,** LPS, Day 7 group: normally inflated alveoli (stars), normal bronchi with peri-bronchial inflammatory cellular infiltration (red arrowheads), mild thick inter-alveolar septa (white arrowheads), mild infiltrated inflammatory cells mostly are alveolar macrophages (thin arrows). **f-i**, Histological score of lung injury in each group, bronchial alteration (**f**), inflammation (**g**), inter-alveolar septal thickness (**h**), total lesion score (**i**). **j**, Flow cytometry counts of neutrophils, macrophages, and B cells in the lung post LPS treatment. **k**, Flow cytometry display of Cyp11a1-mCherry expression in the lung at different time points post LPS treatment. Left panel: representative FACS plot. Right panel: Respective percentage and count of Cyp11a1^+^CD45^+^ cells. **l**, The proportion of immune cell types within Cyp11a1⁺CD45⁺ cells in lung at day 7 post LPS treatment. **m & n,** Reanalyzed single-cell transcriptomics data from lung of LPS-treated mice at different time points post LPS treatment (GSE218884). **m,** Dot plot shows the proportion and average expression of Cyp11a1 across all cells at day 0, day 3, and day 7 post LPS treatment. **n,** dittoBarPlot showing the proportion of different cell types expressing Cyp11a1. Bars indicate the mean ± s.d. The *P* value was calculated using a two-way ANOVA (b) and one-way ANOVA (f-k) with corrections for multiple comparisons. Panel (a) was created with BioRender.

Interestingly, Cyp11a1-expressing immune cells (i.e., steroidogenic immunocytes) were depleted during the inflammatory phase and replenished in the lung during the resolution phase following LPS-induced ALI (Fig. 1k). Both the percentage and absolute number of Cyp11a1⁺CD45⁺ cells were significantly reduced on day 3 but showed a notable increase by day 7 following LPS treatment (Fig. 1k). Basophils were identified as the primary *de novo* steroidogenic immune cells in the lung during the recovery phase following ALI (Fig. 1l). To further validate our findings, we reanalyzed a publicly available single-cell RNA sequencing (scRNA-seq) dataset generated from FACS-sorted CD45^+^ lung cells from an LPS-induced ALI mouse model^39^. Notably, our analysis revealed that *Cyp11a1* expression was markedly reduced during the inflammatory phase (day 3) and elevated during the resolution phase (day 7) (Fig. 1m and Extended Data Fig. 1a), consistent with our findings. scRNA-seq also confirmed that Basophils are the main *de novo* steroidogenic cells during the resolution phase following LPS-induced ALI (Fig. 1n). We also reanalyzed a publicly available scRNA-seq dataset generated from FACS-sorted lung CD45^+^ cells from *Escherichia coli* induced pneumonia model^40^. Consistent with our findings, the analysis revealed that Cyp11a1 expression was markedly reduced on day 3 following bacterial infection and increased by day 7 during the resolution phase (Extended Data Fig. 1b). scRNA-seq analysis further identified basophils as the predominant Cyp11a1-expressing immune cell population in the lung (Extended Data Fig. 1c). Given that ALI can also be induced by viral infections, we reanalyzed bulk RNA sequencing data generated from SARS-CoV-2–infected mouse lungs^41^. Cyp11a1 expression was reduced during the inflammatory phase and significantly upregulated during the resolution phase following virus infection (Extended Data Fig. 1d).

To investigate the role of Cyp11a1-expressing cells in recovery following ALI, we generated mice with immune cell-specific *Cyp11a1* deletion by crossing *Vav1^Cre^* with *Cyp11a1^fl/fl^* mice (Fig. 2a). Compared to control mice, *Cyp11a1^fl/fl^;Vav1^Cre^*(Cyp11a1-cKO) mice failed to resolve the inflammation and recover following LPS-induced ALI. The sickness score (Fig. 2b) indicated that both *Cyp11a1^fl/fl^; Vav1^Cre^* and control mice exhibited a comparable initial inflammatory response; however, only *Cyp11a1^fl/fl^;Vav1^Cre^* mice fail to resolve inflammation and recover. Mice were euthanized on day 5 due to *Cyp11a1^fl/fl^;Vav1^Cre^* mice reaching the humane endpoints. Histological examination of the lungs using H&E staining was performed on day 3 and 5 post LPS-treatment. On day 3, both *Cyp11a1^fl/fl^;Vav1^Cre^*and control mice exhibit comparable pathological changes in the lung (Fig. 2c-h). However, on day 5 post treatment (Fig. 2i-n), we observed a significant difference in bronchial alteration (Fig. 2k), inflammation (Fig. 2l), inter-alveolar septal thickness (Fig. 2m), and total lesion score (Fig. 2n) between *Cyp11a1^fl/fl^;Vav1^Cre^* and control mice, which suggest that while control mice had largely recovered, *Cyp11a1^fl/fl^;Vav1 ^Cre^* mice exhibited signs of unresolved lung inflammation and injury.

**Fig. 2.**
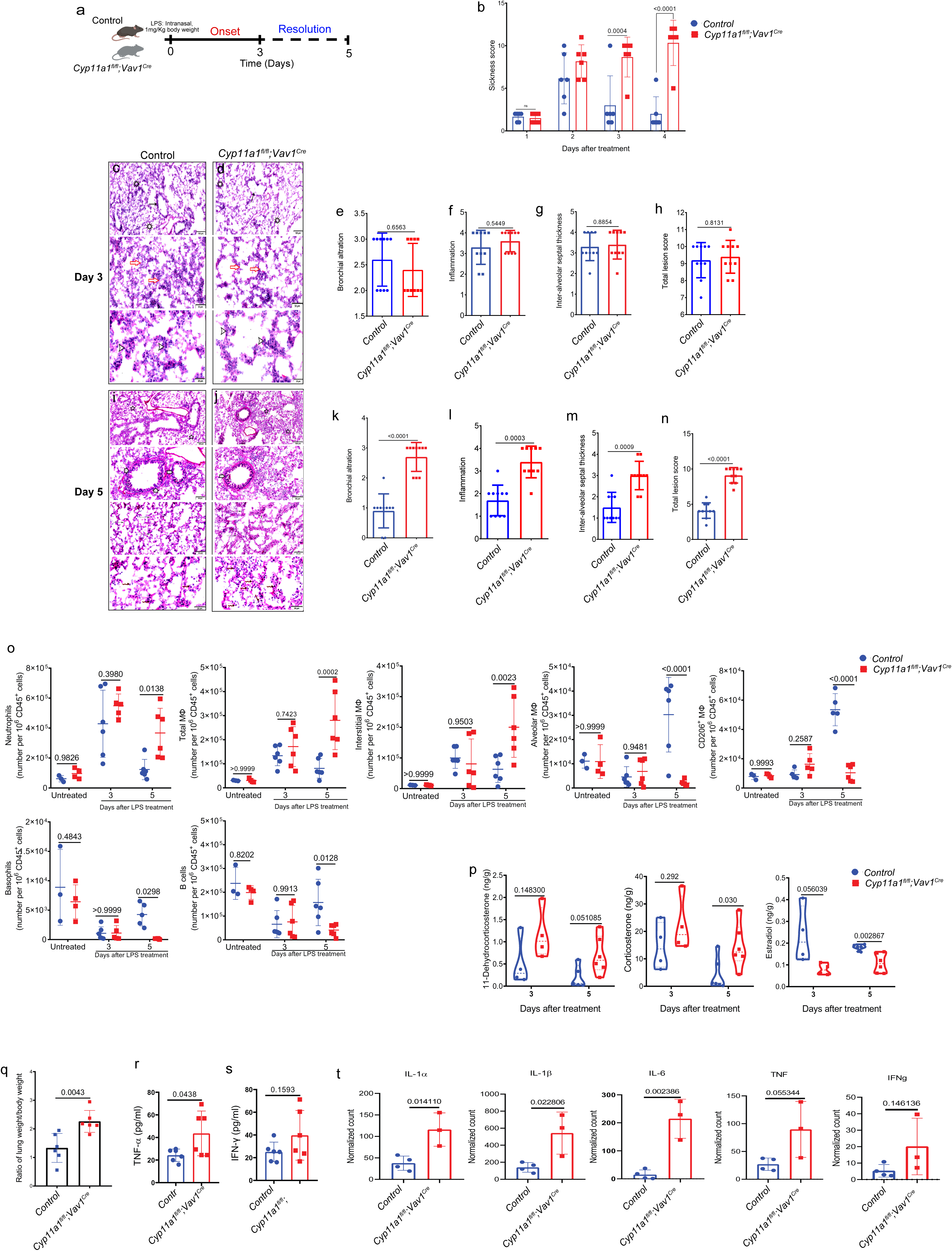
Deletion of Cyp11a1 in immune cells compromises resolution of inflammation and recovery following LPS-induced lung injury. **a,** Schematic representation of LPS-induced ALI experiment in *Cyp11a1^fl/fl^;Vav1^Cre^* and control mice. Mice were anesthetized and intranasally administered 50 µL of LPS. **b**, Sickness score. **c** & **d**, representative images of hematoxylin and eosin (H&E) staining of lung sections from *Cyp11a1^fl/fl^;Vav1^Cr^* (n=6) control 7 (n=6) at day 3 post LPS treatment, alveolar collapse (stars), degenerated bronchioles with peri-bronchial inflammatory cellular infiltration (thin arrows), interstitial pneumonia, thick interalveolar septa with marked inflammatory cellular infiltration (red arrows), and infiltrated inflammatory cells are mostly macrophages (arrowheads). **e-h**, Histological score of lung injury in each group at day 3 post LPS treatment, bronchial alteration (**e**), inflammation (**f**), inter-alveolar septal thickness (**g**), total lesion score (**h**). **i,j**, representative images of Hematoxylin and eosin (H&E) staining of lung sections from *Cyp11a1^fl/fl^;Vav1^Cr^* (n=6) control 7 (n=6) at day 5 post LPS treatment, Control group showed normal inflated alveoli (stars), normal bronchial epithelium with mild peribronchial inflammatory cellular infiltration (arrowheads), normal inter-alveolar septa (thin arrows), and mild infiltrated alveolar macrophages (zigzag arrows) (**i**). *Cyp11a1^fl/fl^;Vav1^Cre^* group showed moderate edematous fluid infiltrated with mononuclear inflammatory cells occluding the alveolar space (stars), epithelium with mild peribronchial inflammatory cellular infiltration (arrowheads), interstitial alveolitis with thick inter-alveolar septa (thin arrows), and marked infiltrated immune cells (Zigzag arrows) (**j**). **k-n**, Histological score of lung injury in each group at day 3 post LPS treatment, bronchial alteration (**k**), inflammation (**l**), inter-alveolar septal thickness (**m**), and total lesion score (**n**). **o,** flow cytometry counts of neutrophils, total MΦs, Ims, Ams, CD206^+^MΦs, DCs, basophils, and B cells in the lung at day 0, 3, and 5 post LPS treatment. **p**, profiling different steroids in the lung at day 3 and 5 post LPS treatment. **q**, ratio of lung weight to body weight at 5 post LPS treatment. **r**, ELISA quantification of TNF-α concentration in the lung at 5 post LPS treatment. **s**, Quantification of IFN-γ concentration in the lung at 5 post LPS treatment using ELISA. **t**, normalized count of the expression of inflammatory cytokines between *Cyp11a1^fl/fl^;Vav1^Cre^*and control mice at day 5 post LPS-treatment detected by bulk RNA-sequencing. Bars indicate the mean ± s.d. The *P* value was calculated using a two-way ANOVA (b,c, and p) and two-tailed unpaired *t*-test (q-t). Panel a created with BioRender.com.

*Cyp11a1^fl/fl^;Vav1^Cre^* and control mice exhibit comparable lung immune cell profiles at steady state (untreated) and on day 3 after LPS treatment (Fig. 2o). However, on day 5 after LPS treatment, significant differences emerged in immune cell populations during the resolution phase. By day 5 after treatment, neutrophil numbers had returned to baseline levels in control mice but remained elevated in *Cyp11a1^fl/fl^;Vav1^Cre^*mice, indicating unresolved inflammation (Fig. 2o). Similarly, the number of total MΦs and IMS remained significantly elevated in *Cyp11a1^fl/fl^;Vav1^Cre^* mice (Fig. 2o). In contrast, we observed a significant reduction in the count of basophils, AMs, CD206^+^ MΦs, and B cells (Fig. 2o) during resolution phase in *Cyp11a1^fl/fl^;Vav1^Cre^* compared to control group. We also observed a marked decrease in T cells number (Extended Data Fig. 2a). There were no significant differences in dendritic (DCs) (Extended Data Fig. 2a), natural killer (NK) (Extended Data Fig. 2a), and gamma delta (γδ) T (Extended Data Fig. 2b) cells between the two groups. We observed a significant decrease in CD4^+^, CD8^+^, and FOXP3^+^ T cells in *Cyp11a1^fl/fl^;Vav1^Cre^* on day 5 post treatment (Extended Data Fig. 2b).

Given that the immune cells in *Cyp11a1^fl/fl^;Vav1^Cre^*mice are deficient in intrinsic *de novo* steroidogenic pathway and cannot produce steroids *de novo*, we sought to profile the steroids present during the inflammation and resolution phases to clarify the contribution and the role of immune cells derived steroids during the resolution phase of ALI. On day 5, we observed a markedly higher level of 11-Dehydrocorticosterone in *Cyp11a1^fl/fl^;Vav1^Cre^* (Fig. 2p). Similarly, corticosterone levels remained significantly elevated in *Cyp11a1^fl/fl^;Vav1^Cre^*mice, indicating a sustained stress response and ongoing disease state (Fig. 2p). In contrast, we observed a marked reduction in estradiol concentration on day 3 in *Cyp11a1^fl/fl^;Vav1^Cre^*, which reached statistical significance by day 5 post-LPS treatment (Fig. 2p). There were no significant differences in the levels of pregnenolone, progesterone, 11-deoxycorticosterone, 17aOH-Progesterone, and testosterone between *Cyp11a1^fl/fl^;Vav1^Cre^*and control mice (Extended Data Fig. 2c). We didn’t observe any significant difference across all tested steroids between the two groups of mice on day 3 post treatment (Fig. 2p and Extended Data Fig. 2c).

To further investigate why *Cyp11a1^fl/fl^;Vav1^Cre^* mice fail to resolve inflammation, we analyzed the key inflammatory cytokines at day 5 following ALI. Notably, on day 5 post-treatment, the lungs of *Cyp11a1^fl/fl^;Vav1^Cre^*mice appeared markedly swollen, and lung-to-body weight ratios (Fig. 2q) were significantly higher than those of control mice, indicating unresolved inflammation. Similarly, the percentage of iNOS-expressing MΦs was significantly higher in *Cyp11a1^fl/fl^;Vav1^Cre^*mice compared to control (Extended Data Fig. 2d). *Cyp11a1^fl/fl^;Vav1^Cre^*mice also exhibited a significantly higher concentration of TNF-α in the lung tissue compared to control (Fig. 2r). Compared to neutrophils, DCs, T, B, and NK cells, only MΦs exhibited a significant upregulation of TNF-α (Extended Data Fig. 2e). We didn’t observe any significant difference in the overall concentration of IFN-γ between the two groups of mice (Fig. 2s), however MΦs from *Cyp11a1^fl/fl^;Vav1^Cre^* showed a significantly higher expression of IFN-γ compared to other immune cells (Extended Data Fig. 2f). To further validate cytokine dysregulation during the resolution phase of LPS-induced acute lung injury (ALI), we assessed the transcriptomes of key inflammatory cytokines using bulk RNA-sequencing analysis. By day 5 post-treatment, *Cyp11a1^fl/fl^;Vav1^Cre^* mice exhibited significantly elevated transcript levels of IL-1α, IL-1β, IL-6, and TNF compared to controls (Fig. 2t). However, no significant differences in the levels of these cytokines were observed at steady state (Extended Data Fig. 2g) or on day 3 (Extended Data Fig. 2h). Collectively, these results suggest that while *Cyp11a1^fl/fl^;Vav1^Cre^*mice mount an immune response comparable to controls, they fail to resolve inflammation and recover effectively.

To investigate the genome-wide transcriptomic differences between *Cyp11a1^fl/fl^;Vav1^Cre^* and control mice following LPS-treatment, we performed bulk RNA-seq of lungs from both groups of mice. On day 3 post-ALI, only a few genes were significantly dysregulated; however, by day 5, there was a substantial increase in transcriptional alterations (Fig. 3a-d). While both groups showed comparable expression of inflammatory genes on day 3, by day 5, these genes had resolved in control mice but remained elevated in *Cyp11a1^fl/fl^;Vav1^Cre^* mice (Fig. 3e), indicating persistent inflammation. In contrast, genes involved in the resolution of inflammation and tissue repair were downregulated in *Cyp11a1^fl/fl^;Vav1^Cre^* mice compared to control (Fig. 3f). To further examine gene expression dynamics during resolution, differentially expressed genes between days 3 and 5 within each group were used to cluster the samples. Day 3 control samples clustered together with both day 3 and day 5 samples from *Cyp11a1^fl/fl^;Vav1^Cre^* mice, suggesting a shared inflammatory profile. Notably, day 5 control samples exhibited a distinct expression profile and formed a separate cluster, reflecting a resolved transcriptional signature (Fig. 3g). Pathway enrichment analysis revealed widespread dysregulation in *Cyp11a1^fl/fl^;Vav1^Cre^*mice, with prominent upregulation of pathways linked to inflammatory responses, innate immune response, and acute phase response (Fig. 3h,i).

**Fig. 3.**
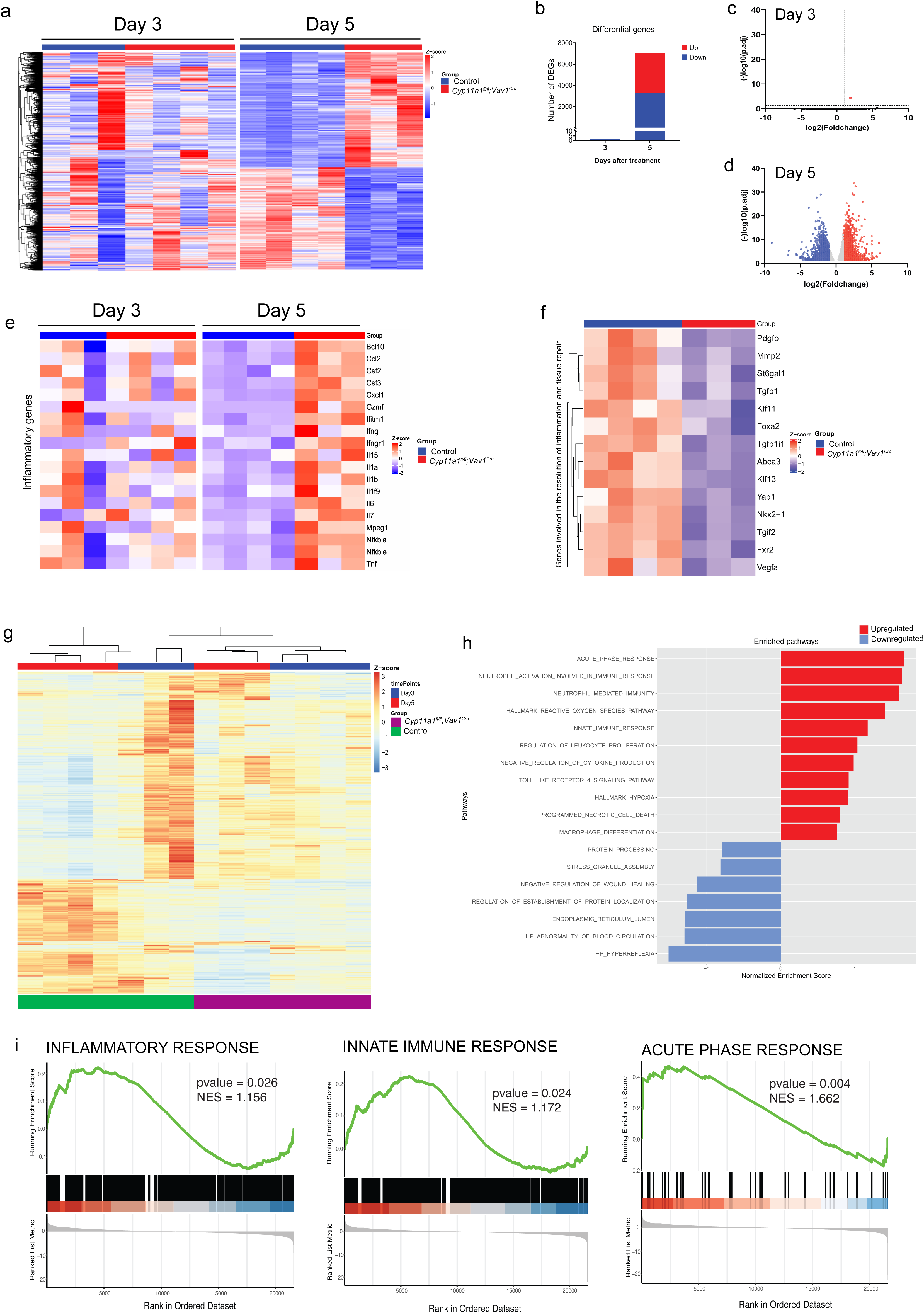
Deletion of *Cyp11a1* in immune cells compromises resolution of inflammation response following LPS-induce lung injury. **a**, Heatmap showing normalized gene expression counts across samples at days 3 and 5 post LPS-treatment. In heatmap red: upregulation, in blue: down-regulation showing the differentially expressed genes. **b**, number of differentially expressed genes at days 3 and 5 post LPS-treatment. **c**,**d**, Volcano plot showing differentially expressed genes in control and *Cyp11a1^fl/fl^;Vav1^Cre^* mice at day 3 and 5 post LPS-treatment. Each dot represents an individual gene. Red dots indicate significantly upregulated genes in *Cyp11a1^fl/fl^;Vav1^Cre^* mice relative to control, blue dots indicate significantly downregulated genes in *Cyp11a1^fl/fl^;Vav1^Cre^*, and grey dots represent genes that did not meet the significance threshold. **e**, heatmap of differentially expressed inflammatory genes in in control and *Cyp11a1^fl/fl^;Vav1^Cre^*mice at days 3 and 5 post LPS-treatment. **f,** heatmap of differentially expressed genes involved in the resolution of inflammation and tissue repair in control and *Cyp11a1^fl/fl^;Vav1^Cre^* mice at day 5 post LPS-treatment. **g**, differentially expressed genes between days 3 and 5 within control and *Cyp11a1^fl/fl^;Vav1^Cre^* mice were used to cluster the samples at days 3 and 5 post LPS-treatment. **h**, the hallmark pathways upregulated (red) and downregulated (blue) in control and *Cyp11a1^fl/fl^;Vav1^Cre^* mice at day 5 post LPS-treatment. **i,** upregulated pathways in control and *Cyp11a1^fl/fl^;Vav1^Cre^* mice at day 5 post LPS-treatment, the GSEA enrichment plots show the upregulated inflammatory response, innate immune response, and acute phase response in *Cyp11a1^fl/fl^;Vav1^Cre^* mice.

Resolution of inflammation is essential for effective lung repair following injury, whereas persistent inflammation leads to a dysregulated repair response and fibrosis^42, 43, 44, 45, 46, 47, 48, 49^. Given that *Cyp11a1^fl/fl^;Vav1^Cre^*mice failed to resolve inflammation compared to controls, we investigated whether Cyp11a1 deficiency in immune cells impairs lung repair and promote fibrosis in the context of LPS-induced acute lung injury (ALI). To prolong the survival of *Cyp11a1^fl/fl^;Vav1^Cre^* mice and enable the study of lung repair and fibrosis, we reduced the LPS dose to 0.5 mg/kg of body weight (Fig. 4a). Regardless of the milder LPS dose, we still observe significant differences in the sickness score between *Cyp11a1^fl/fl^;Vav1^Cre^*and control mice during the resolution phase following LPS administration (Fig. 4b). On day 14 post LPS-treatment, we did not observe any significant differences in the lung immune profiles, except for AMs and basophils, which were markedly reduced in *Cyp11a1^fl/fl^;Vav1^Cre^* mice compared to controls (Fig. 4c and Extended Data Fig. 3a,b). Similarly, there were no significant changes in the expression of TNF-α (Extended Data Fig. 3c), IFN-γ (Extended Data Fig. 3d), and iNOS^+^ MΦs (Extended Data Fig. 3e). Lung tissue histology was examined on day 14 post LPS treatment using H&E and Masson’s trichrome staining. Both H&E and Masson’s trichrome staining revealed substantial fibrotic areas and collagen deposition in the lung of *Cyp11a1^fl/fl^;Vav1^Cre^* compared to control (Fig. 4d-j). Lungs from control mice showed near complete resolution of inflammation and return of the airways and alveolar compartments into near-normal histology (Fig. 4f,). In contrast *Cyp11a1^fl/fl^;Vav1^Cre^*lungs showed focal areas of fibrosis, bronchial epithelial cells with foamy cytoplasm, and significantly higher fibrosis score (Fig. 4e,f). There was no significant difference in the inflammation score between the two groups of mice (Fig. 4g). Masson’s trichrome staining indicated significant increase in collagen deposition in *Cyp11a1^fl/fl^;Vav1^Cre^* lungs compared to control (Fig. 4h-j). Furthermore, quantification of α-smooth muscle actin (α-SMA)-positive cells in the alveolar region revealed a significant increase in the abundance of myofibroblast (Fig. 4k, l), which is the hallmark of pulmonary fibrosis. The accumulation of these α-SMA^+^ myofibroblast suggests a progression toward a fibrotic phenotype in the lung tissue of *Cyp11a1^fl/fl^;Vav1^Cre^* mice. These results suggest that a defective inflammation-resolution mechanism in Cyp11a1-KO mice may contribute to the development of lung fibrosis, a condition commonly observed in severe respiratory infections^42, 50, 51, 52, 53, 54, 55, 56^.

**Fig. 4.**
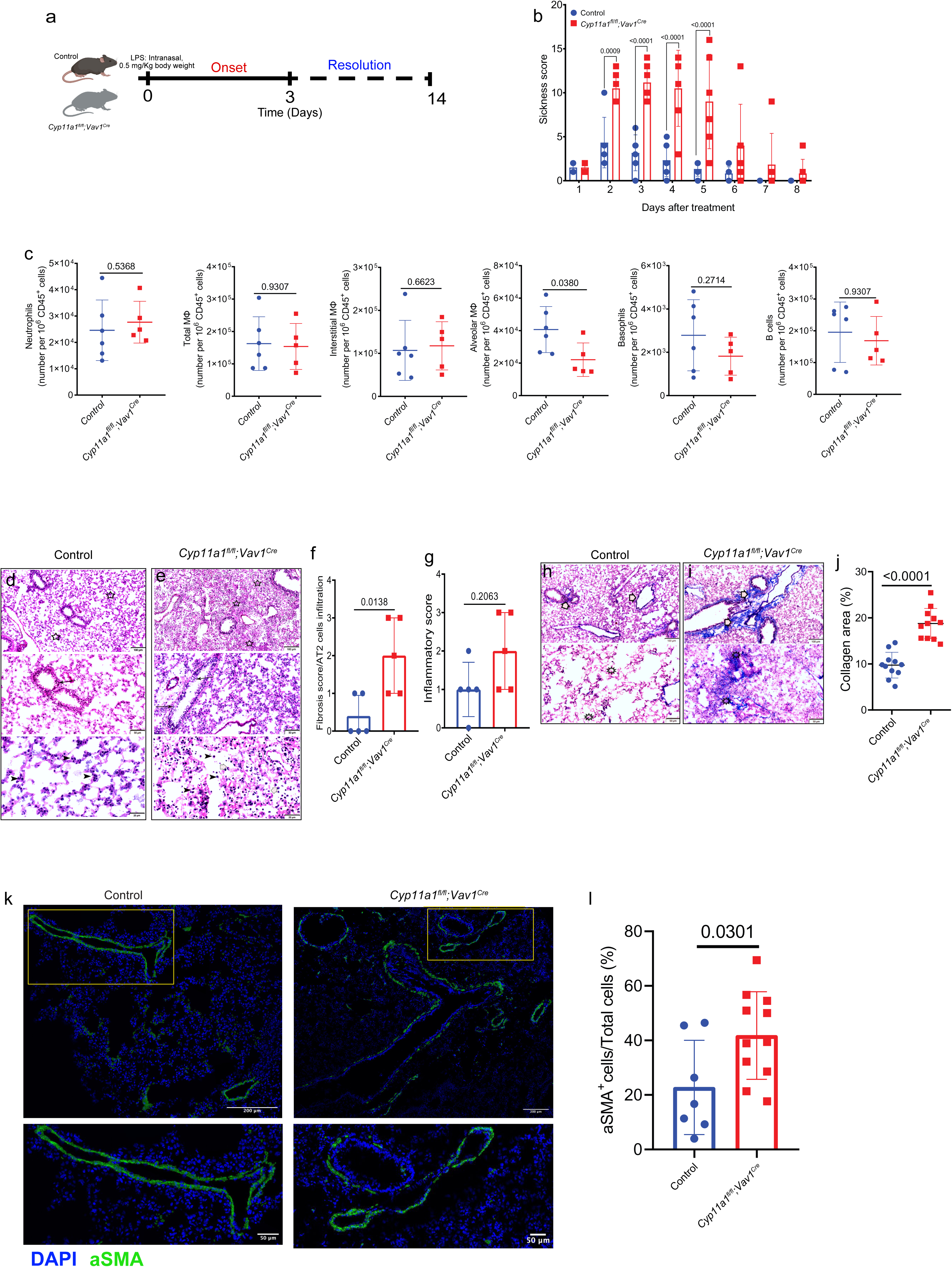
Deletion of Cyp11a1 in immune cells compromises lung repair and promote tissue fibrosis following LPS-induce lung injury. **a,** Schematic representation of LPS-induced ALI experiment in *Cyp11a1^fl/fl^;Vav1^Cre^*and control mice. **b**, sickness score. **c**, flow cytometry counts of neutrophils, total MΦs, Ims, Ams, CD206^+^MΦs, DCs, basophils, and B cells in the lung at day 14 post LPS treatment **d,e**, representative images of hematoxylin and eosin (H&E) staining of lung sections from *Cyp11a1^fl/fl^;Vav1^Cre^* (n=6) control 7 (n=6) at day 14 post LPS treatment. Control group showed normal inflated alveoli (stars) without evidence of fibrosis, normal bronchioles (arrow), and normally infiltrated alveolar macrophages (black arrowheads) (f). *Cyp11a1^fl/fl^;Vav1^Cre^*group showed fibrotic areas with intense cellular proliferation in the lung parenchyma (stars), bronchiolitis with foamy bronchial epithelial cells (arrows), inflammatory cellular infiltration mostly alveolar macrophages (black arrowheads), and neutrophils (white arrows) (g). **f**,**g**, histological score of lung fibrosis and inflammation in each group at day 14 post LPS treatment, Fibrosis score/AT2 cells infiltration (f) and inflammation (g). **h**,**i** representative images of lung sections from *Cyp11a1^fl/fl^;Vav1^Cr^*(n=6) control 7 (n=6) at day 14 post LPS treatment stained with Masson’s trichrome. Control group showed normal fine collagenous fibrous threads around bronchioles (arrowheads), and no evidence of fibrosis in lung parenchyma (stars) (h). *Cyp11a1^fl/fl^;Vav1^Cre^*group showed fibrotic areas around bronchioles (arrowheads) and focally associated with intense cellular proliferation in the lung parenchyma (stars) (i). **j**, quantification of collagen area from Masson’s trichrome stained lung tissue. **k**,**l**, immunostaining images and quantification of aSMA positive cells from lungs of *Cyp11a1^fl/fl^;Vav1^Cr^*and control mice at day 14 post LPS treatment. Bars indicate the mean ± s.d. The *P* value was calculated using a two-way ANOVA (b) and two-tailed unpaired *t*-test (c-l). Panel a created with BioRender.com.

Given that the basophils were identified as the main *de novo* steroidogenic immune cells in the lung, we used *Cpa3*-derived Cre to knock out *Cyp11a1* in basophils^57^. We administered a mild dose (0.1mg/Kg of body weight) of LPS to *Cyp11a1^fl/fl^;Vav1^Cre^*, *Cyp11a1^fl/fl^;Cpa3^Cre^*, and control mice to assess their response under mild inflammatory conditions (Fig. 5a). Interestingly, both *Cyp11a1^fl/fl^;Vav1^Cre^*and *Cyp11a1^fl/fl^;Cpa3^Cre^* mice exhibited significantly higher sickness scores compared to controls (Fig. 5b). However, there was no significant difference in sickness scores between the two Cyp11a1-knockout groups. H&E staining revealed that control mice displayed normal parenchyma, intact bronchiolar epithelium, and minimal inflammatory infiltration (Fig. 5c). In contrast, *Cyp11a1^fl/fl^;Vav1^Cre^*(Fig. 5d) and *Cyp11a1^fl/fl^;Cpa3^Cre^*(Fig. 5e) mice exhibited moderate to heavy parenchymal infiltration with *Cyp11a1^fl/fl^;Vav1^Cre^*showing obliterated alveolar spaces and *Cyp11a1^fl/fl^;Cpa3^Cre^*displaying mild inflammation in terminal bronchioles and interalveolar walls. Total inflammatory score was significantly higher in the two Cyp11a1-KO groups compared to control (Fig. 5f).

**Fig. 5.**
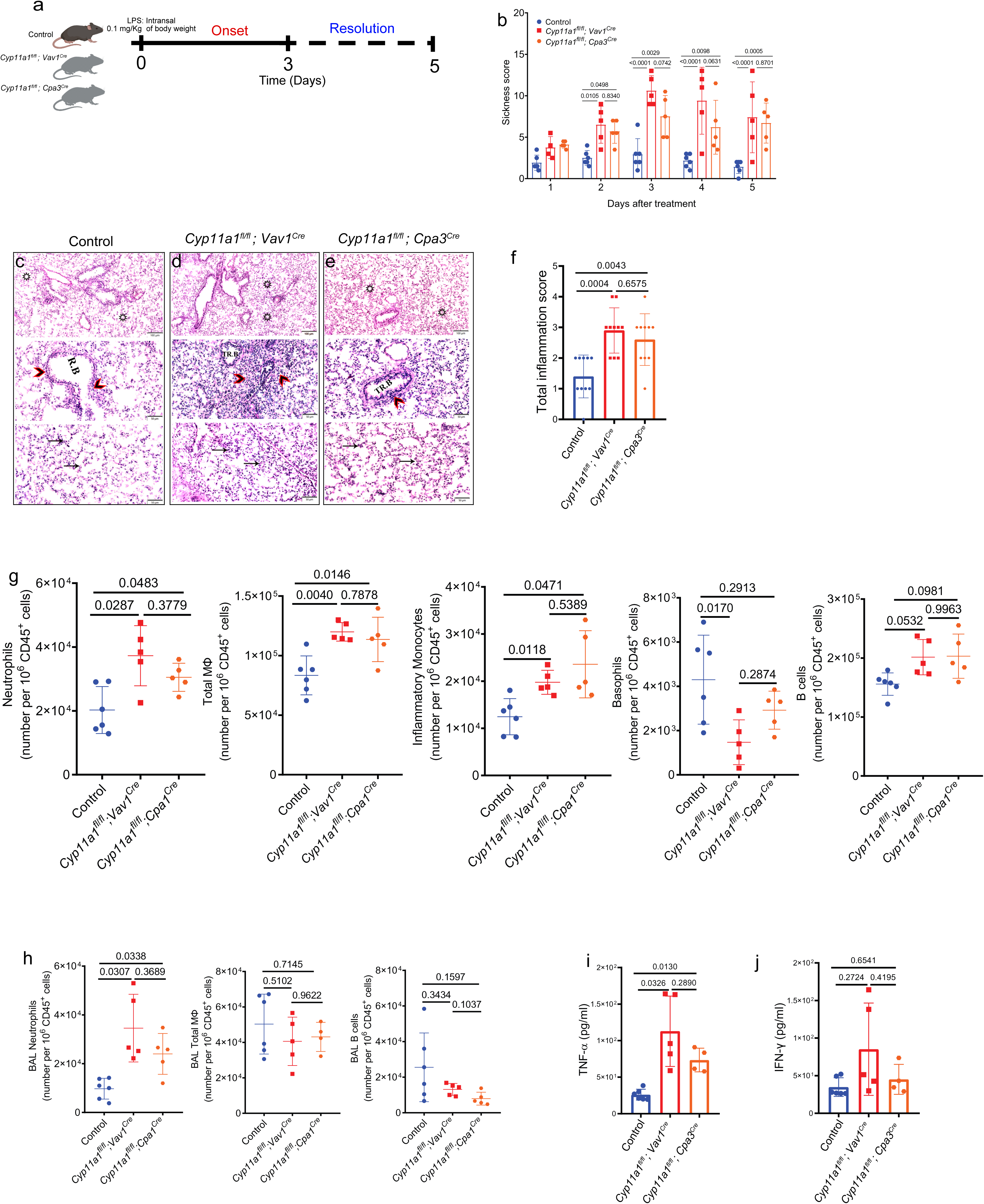
Deletion of Cyp11a1 in basophils compromises resolution of inflammation following LPS-induce lung injury. **a,** Schematic representation of LPS-induced ALI experiment in *Cyp11a1^fl/fl^;Vav1^Cre^*, *Cyp11a1^fl/fl^;Cpa3^Cre^*, and control mice. **b**, thickness score. **c-e**, representative images of Hematoxylin and eosin (H&E) staining of lung sections from *Cyp11a1^fl/fl^;Vav1^Cre^* (n=5), *Cyp11a1^fl/fl^;Cpa3^Cre^*, and control 7 (n=6). **c**, control group exhibited a normal aerated parenchymal tissue (stars), respiratory bronchioles (R.B) with intact lining epithelium and mild inflammatory cells infiltration (arrowheads), normal alveoli with more or less normal interstitial cellular infiltration (arrows). **d**, *Cyp11a1^fl/fl^;Vav1^Cre^* showed moderate cellular infiltration in the aerated parenchyma (stars) and heavy inflammatory cellular infiltration obliterated focal aerated alveolar spaces (arrowheads), interstitial inflammatory cellular infiltration (arrows). **e**, *Cyp11a1^fl/fl^;Cpa3^Cre^* showed moderate cellular infiltration in the aerated parenchyma (stars), terminal bronchiole (TR.B) showed mild inflammatory cell infiltration (arrowhead), and mild proliferating inflammatory cells in the interalveolar wall (arrows). **f**, histological score of lung inflammation in each group at day 5 post LPS treatment. **g**, flow cytometry counts of neutrophils, total MΦs, inflammatory monocytes, basophils, and B cells in the lung at day 5 post LPS treatment. **h**, flow cytometry counts of Neutrophils, MΦs, T, and B cells in BAL fluid at day 5 post LPS treatment. **i**,**j,** ELISA quantification of the concentration of TNF-α (i) and IFN-γ (**j**) in BALF at day 5 post LPS treatment. Bars indicate the mean ± s.d. The *P* value was calculated using a two-way ANOVA (b) and one-way ANOVA (f-j). Panel a created with BioRender.com.

Compared to control, both *Cyp11a1^fl/fl^;Vav1^Cre^* and *Cyp11a1^fl/fl^;Cpa3^Cre^* mice exhibited a significant increase in neutrophils, macrophages (MΦs), inflammatory monocytes, and B cells in the lung (Fig. 5g) as well as Interstitial and Alveolar macrophages (Extended Data Fig. 4a). In contrast, basophil counts were markedly reduced in the two Cyp11a1-KO groups, particularly in the *Cyp11a1^fl/fl^;Vav1^Cre^*group compared to control (Fig. 5g). However, no significant changes were observed in dendritic cells, T cells, NK cells, or mast cells populations across the treatment groups (Extended Data Fig. 4b). In fact, there were no significant differences in the count of T cells subsets across the treatment groups (Extended Data Fig. 4c). TNF-α expression was significantly upregulated in neutrophils, MΦ, DCs, T, and B cells in both *Cyp11a1*-KO groups compared with controls (Extended Data Fig. 4d). Notably, Inflammatory monocytes and NK cells did not show a significant difference in TNF-α expression (Extended Data Fig. 4d). In contrast to TNF-α, IFN-γ levels did not differ significantly across immune cells in the various treatment groups (Extended Data Fig. 4e).

To further elucidate the impact of Cyp11a1 deficiency in basophils on resolution of lung inflammation, we analyzed the immune cell composition and cytokine levels in bronchoalveolar lavage (BAL) fluid across different treatment groups on Day 5 following LPS administration. Notably, neutrophil counts were significantly higher in BAL collected from both *Cyp11a1^fl/fl^;Vav1^Cre^*and *Cyp11a1^fl/fl^;Cpa3^Cre^* groups compared to controls (Fig. 5h). In contrast, no significant differences were observed in BAL MΦ, B cell, or T cell numbers among the groups (Fig. 5h and Extended Data Fig. 4f). A significant difference was observed in TNF-α levels in the BAL of the Cyp11a1-KO groups compared with controls (Fig. 5i). In contrast, IFN-γ concentrations did not differ significantly across the treatment groups (Fig. 5j). These findings indicate that basophil-derived *de novo* steroids play a critical role in resolving inflammation and promoting recovery following ALI.

In summary, our findings establish a critical role for *de novo* steroidogenesis by immune cells, particularly by basophils, in orchestrating the resolution of inflammation and facilitating lung repair following acute lung injury (ALI). Our study revealed that Cyp11a1-expressing immune cells are dynamically regulated during the inflammatory and resolution phases. These cells, especially basophils, are depleted during the peak of inflammation yet enriched during the resolution phase, coinciding with recovery of lung function and immune homeostasis. Mice with immune cell-specific deletion of *Cyp11a1* fail to resolve inflammation, as evidenced by sustained inflammatory cells and persistent pro-inflammatory cytokines. These mice exhibit persistent lung injury, exaggerated fibrotic remodeling, and increased accumulation of α-SMA^+^ myofibroblast, ultimately leading to defective tissue repair and fibrosis. Notably, deletion of *Cyp11a1* specifically in basophils and mast cells similarly impaired resolution and recapitulated key aspects of the phenotype observed in the global immune cell *Cyp11a1* deletion, underscoring the functional relevance of basophil as the main *de novo* steroidogenic immune cells in the lung. These findings may open new avenues for therapeutic interventions aimed at enhancing intrinsic *de novo* steroidogenesis in immune cells to promote resolution of lung inflammation and prevent progression to fibrosis in respiratory diseases.

While our study provides compelling evidence for the role of immune cell-intrinsic steroidogenesis in resolving acute lung injury, downstream mechanisms and molecular targets of immune-derived steroids were not fully elucidated. The study focused on histologic and immunologic parameters of resolution but did not assess long-term pulmonary function, which would provide clinically relevant insight into tissue recovery. In the future, further validation in other models, including infectious or mechanical injury, would strengthen the translational relevance of these findings.

## Methods

### Mice

All animal procedures were conducted in accordance with the UK Animals (Scientific Procedures) Act 1986 and its Amendment Regulations 2012, adhering to the UK Animals in Science Regulation Unit’s Code of Practice for the Housing and Care of Animals Bred, Supplied or Used for Scientific Purposes. Experiments were performed under UK Home Office Project Licence PPL (PP3952258 and PP4938782) and approved by the institutional Animal Welfare and Ethical Review Body. Sample sizes were determined based on prior experimental experience and a priori power analysis using G*Power software. Mice were housed in a specific pathogen-free facility under a 12-hour light/dark cycle. Genotyping was performed by Transnetyx. Cyp11a1-mCherry reporter and *Cyp11a1^fl/fl^*mice were generated by the Sanger Institute as described previously ^19^. Immune cell-specific Knockout mice (*Cyp11a1^fl/fl^;Vav1^Cre^*) and Mast cell and basophil-specific knockout mice (*Cyp11a1^fl/fl^;Cpa3^Cre^*) were generated by crossing *Cyp11a1^fl/fl^* mice with *Vav1*-Cre and Cpa3-Cre mice (Jackson Laboratory), respectively. Mice aged 8–16 weeks were used in this study.

### *In vivo* LPS challenge

Mice were anesthetized with Isoflurane and intranasally administered 50 µL of either sterile phosphate-buffered saline (PBS) or lipopolysaccharide (LPS) from Escherichia coli O111:B4 (Sigma-Aldrich) in PBS. LPS was administered based on the body weight of the mouse. The sickness of animals was assessed using a scoring system detailed in Supplementary Table 1 by two independent scientists. Mice were sacrificed at different time points following LPS treatment (3, 5 and 7 days, n = 6 per time point).

### BAL fluid and lung collection and processing

At desired time points post LPS treatment, mice were anesthetized and subsequently, the lungs were lavaged twice with sterile PBS supplemented with 2mM EDTA to collect the BAL Fluid. Following BAL collection, lungs were perfused by cold PBS and harvested into cold PBS supplemented with 2.5% FBS and 2mM EDTA and stored on ice until further processing. BAL was centrifuged and the cell pellet resuspended and used for flow cytometry staining. The cell-free BAL was stored at -80 °C for further analysis. To extract lung leukocytes, lung tissue was initially chopped very finely and then incubated in digestion buffer (DMEM/F12 supplemented with 1mg/ml collagenase A (Roche), 1mg/ml collagenase D (Roche), 35µg/ml mg/ml DNase I (Sigma Aldrich), and 10% FBS) at 37 °C in an agitating mixer (Eppendorf ThermoMixer C) for 30 minutes. The digested tissue was then filtered through 70μm cell strainers (Falcon). Erythrocytes were lysed using 1X RBC lysis buffer (eBioscience) prior to cell counting using a Countess Automated Cell Counter (ThermoFisher).

### Flow cytometry

Cells were stained for viability using either Live dead Ghost dye (CYTEK) or Fixable Near-IR (ThermoFisher) and blocked using purified rat anti-mouse CD16/CD32 purchased from eBioscience, BioLegend, CYTEK, or BD. Surface staining was performed in FACS buffer (PBS with 2.5% FBS and 2mM EDTA) at 4°C for 2 hours in the dark. Cells were washed with FACS buffer and fixed in IC Fixation Buffer (eBioscience) and washed twice in FACS buffer. For experiments involving intranuclear staining, cells were fixed and permeabilized using Foxp3/Transcription factor staining Buffer Kit (eBioscience) for 30 minutes. Subsequently, cells were washed twice with permeabilization buffer (eBioscience) and stained overnight with antibodies in permeabilization buffer(eBioscience). At the end of the incubation, cells were washed with FACS buffer and 5000 Precision Count beads (BioLegend) added per sample. The antibody cocktails used for flow cytometry staining can be found in Supplementary Table 2. Data were acquired on an Aurora Spectral Analyzer (Cytek). Gating strategy used for the quantification of immune cells are provided in Extended Data Fig. 5a–c.

### Bulk RNA sequencing data analysis

RNA was extracted from lung tissue using RNeasy Mini Kit or RNeasy Micro Kit (QIAGEN) following the manufacturer’s protocol. RNA-Seq was performed using high-quality total RNA samples. cDNA libraries were constructed and sequenced using paired-end reads on an Illumina NovaSeq X Plus Series (PE150), yielding an average of 50 million reads for each library. The quality of the raw data was checked with FastQC before and after trimming of FASTQ files with Fastp^58^. We used Hisat2^59^ with default settings for sequence alignment, using Grcm38^60^ as reference genome. Following alignment, duplicated reads were filtered from BAM files using samtools^61^. The GTF file mus_musculus.grcm38.80.gtf was used for gene annotation during alignment^62^. Reads mapped to each gene were enumerated using LiBiNorm count^63^. Further analysis was performed in R version 4.3.2 using DESeq2^64^ and custom scripts. PCA on the normalized read counts showed two outlying samples, which were removed prior to testing for differential expression. Differentially expressed genes (cKO vs control) of Day 5 samples with p-adjusted value < 0.05 were subjected to further analysis regarding gene ontology (GO) and Kyoto Encyclopedia of Genes and Genome (KEGG) pathways. GO enrichment analysis was carried out in R using the topGO package and statistical significances of enrichment were assessed by Fisher’s exact tests and the classic algorithm. KEGG pathway enrichment analysis was performed using the clusterProfiler R package^65^. Significant gene symbols were converted to Entrez Gene IDs and then enrichment analysis was performed against the KEGG Mus musculus database^66^. Enriched KEGG pathways were identified and filtered using a threshold p value < 0.05. The top 15 enriched KEGG pathways were selected for dot plotting. Differentially expressed genes of the Day 5 group by comparing cKO to control samples were used for gene set enrichment analysis (GSEA), which was performed using clusterProfiler R package again. Gene sets associated with ‘hallmarks of inflammatory response’ and immune signatures of neutrophils according to the MSigDB database from the msigdbr package^67^ were tested for enrichment, followed by filtering of results for p values < 0.05, and visualization in graphical formats. Gene symbols were converted to Entrez Gene IDs based on the org.Mm.eg.db annotation database^68^ in all three analyses (GO, KEGG, GSEA). The previously published and publicly available RNA-seq datasets (GSE248773^41^) were analysed for Cyp11a1 expression following SARS-CoV2 infection in mice. This dataset was obtained from the National Center for Biotechnology Information (NCBI) Gene Expression Omnibus (https://www.ncbi.nlm.nih.gov/geo/)

### Single-cell RNA Sequencing, clustering and visualization

The previously published and publicly available scRNAseq datasets used in this study were GSE218884^39^ and PRJEB52332^40^. These datasets were obtained from NBCI and The European Nucleotide Archive (https://www.ebi.ac.uk/ena/browser/view/PRJEB52332), respectively. Data processing followed the standard scRNA-seq integration workflow implemented in Seurat (version 4.1.3)^69^. For clustering, the top 2,000 most variable genes were selected using Seurat’s FindVariableFeatures function. Principal component analysis (PCA) was performed, and Uniform Manifold Approximation and Projection (UMAP) was used to reduce the data to two dimensions for visualization. Cell clustering in PCA space employed the Shared Nearest Neighbor (SNN) approach via Seurat V3’s FindNeighbors and FindClusters functions. The resulting clusters were visualized in UMAP space using the DimPlot function and annotated according to metadata. Gene expression patterns were visualized using the ggplot2 package (v3.5.2) and dittoSeq (v1.2.4). Scaling was applied automatically based on the default settings of these visualization tools^70^.

### Steroid quantification via liquid chromatography mass spectrometry (LC-MS/MS)

Steroid levels in lung tissues were measured using LC-MS/MS as previously described^24^. Briefly, Lung tissue (∼ 50mg) was placed in 2 mL reinforced tubes containing 1.4 mm ceramic beads (FisherScientific). 1 mL of acetonitrile with 0.1% formic acid (v/v) and 20 µL of an isotopically labelled steroid standard mixture were added to the tube. Samples were homogenized at 1 m/s for 30 seconds for 3 cycles using a Bead Ruptor 24 Elite (Omni International) equipped with a CryoCool unit. Supernatants were transferred to a Filter+ plate (Biotage, Sweden), and eluate collected into clean 96-well plates. The filtered homogenate underwent further processing through phospholipid depletion (PLD+) plates (Biotage, Sweden). Eluate was dried, reconstituted in water/methanol (70:30 v/v), and sealed with zone-free plate seals, ready for LC-MS/MS analysis. An I-Class UPLC (Waters, UK) on a Kinetex C18 column (150 × 2.1 mm, 2.6 μm) was used for liquid chromatography with a mobile phase of water and methanol, both containing 0.05 mM ammonium fluoride, starting at 50% methanol (B), increasing to 95%, then returning to 50%. The flow rate was 0.3 mL/min. The column was maintained at 50°C and the autosampler at 10°C, with a 20 µL injection volume. Each analytical run lasted 16 minutes per sample. Steroid detection was performed using a QTRAP 6500+ mass spectrometer (AB Sciex, Warrington, UK) equipped with an electrospray ionization Turbo V ion source. The ion spray voltage was set at +5500 V for positive mode and -4500 V for negative mode, with a source temperature of 600°C. Multiple reaction monitoring parameters were optimized for each steroid, including pregnenolone (P5) with transitions at m/z 317.1→281.1 and 317.1→159.0, and its labeled standard (13C2,d2-P5) at 321.2→285.2. Parameters such as declustering potential (DP), collision energy (CE), and collision cell exit potential (CXP) were set accordingly, with retention times around 10.4 minutes. P5 concentrations were quantified by calculating the ratio of analyte (P5) to its labelled standard (13C2,d2-P5) peak areas, followed by linear regression analysis using MultiQuant 3.0.3 (AB Sciex, UK). This approach was applied to other steroids including aldosterone, progesterone, 17β-estradiol, 5α-dihydrotestosterone, and testosterone.

### Real-time qPCR (RT-qPCR)

To generate complementary DNA (cDNA), 500 ng of total RNA was reverse transcribed into complementary DNA (cDNA) using the High-Capacity cDNA Reverse Transcription Kit (Applied Biosystems, Vilnius, Lithuania). cDNA samples were diluted and subsequently subjected to RT-qPCR assay using Fast SYBR Green Master Mix (Applied Biosystems, Vilnius, Lithuania) following the manufacturer’s instructions. Expression of the cytokine genes was detected using cytokine-specific primers (Supplementary Table 3). GAPDH was used as the endogenous control for normalization.

### Measurement of inflammatory cytokines

TNF-α and IFN-γ cytokines in the homogenized lung tissue and cell-free BAL were quantified using the commercially available mouse enzyme-linked immunosorbent assay (ELISA) kits (eBioscience) according to the manufacturer’s instructions. Lung homogenates were prepared in ice-cold RIPA buffer supplemented with a protease inhibitor cocktail (Roche) using a BEAD RUPTOR 12 tissue homogenizer (OMNI international) followed by centrifugation at ∼ 14,000 x g for 15 minutes at 4°C. Total protein was quantified using the standard BCA protein Assay Kit (Thermo Fisher).

### Immunofluorescence staining, imaging, and quantification

Frozen tissue sections (5um) were fixed in 10% formalin solution (Sigma, HT5011) and permeabilized using 0.2% triton-x. Next, sections were stained with an antibody specific to αSMA (1:200, Proteintech). Sections were then stained with a secondary antibody (1:200, Proteintech). Images were acquired using a Leica SP8 confocal at 20x magnification. Immunofluorescence images were analyzed using ImageJ2 (v2.14). Prior to analysis, all images were background-subtracted and converted to 8-bit grayscale. A manual intensity threshold was applied (set between 5 and 255) to isolate fluorescently labelled cells from background signals. Images were then converted to binary masks, with foreground pixels representing positive signal and background pixels set to zero. To resolve adjacent or clustered cells, a watershed algorithm was applied to the binary mask to ensure accurate object separation. Quantification was performed using the “Analyze Particles” function with a minimum size threshold of 50 pixels to exclude background noise or artifacts. Quantitative data was generated by first counting total cells in alveolar sections as those stained with DAPI and then followed by quantification of aSMA+ cells. The percentage of aSMA+ cells was calculated as total (aSMA+/DAPI+)*100.

### Histopathological examination, quantification, and scoring

Frozen lung tissue sections (5um) were dried, fixed in precooled acetone solution, and stained with hematoxylin and eosin (H&E) following the standard methodology^71^. Sections were imaged and scanned by bright-field microscopy using the high-throughput slide-scanner NanoZoomer system (v2.0, Hamamatsu) with a 20× objective and analyzed using Nanozoomer digital pathology software. Each animal was scored according to the recorded histopathological examination^72^. A visual field inspection of at least 10 tissue specimen sections from each experimental group was performed to record the semiquantitative scoring of histopathological changes. Tissue alterations were scored according to the following criteria: “0”—none, “1” “— <25% “2”—26–50% “3”—51 — 75%, and “4” >75% respectively). The scores were then added to create a final score ranging from 0 to 16 ^73, 74^. Lung tissue sections were scored according to alterations in bronchioles, inflammation and edema, inflation changes, and interalveolar septal thickness. All according to the nature and extent of the lesion and its occurrence frequency in randomly selected tissue sites ^74^. The total lesion score of the last experiment was scored according to alterations in bronchioles, the severity of inflammation “0”—none, “1” “— <25% “2”—26–50% “3”—51 — 75%, and “4” >75% respectively), and then the scores were added to create a final score ranging from 0 to 8.

Lung sections were stained with Masson’s trichrome stain, where the blue color indicates collagen deposition and fibrosis. Tissue sections were fixed in 4% paraformaldehyde for 1 hour at room temperature, followed by overnight incubation in Bouin’s solution. After washing, sections were stained with Weigert’s iron hematoxylin, followed by blueing under warm tap water. Subsequently, Cytoplasm and muscle fibers were stained red using Biebrich scarlet-acid fuchsin for 5 minutes. Sections were then washed and treated with phosphotungstic/phosphomolybdic acid, followed by incubation with aniline blue to visualize collagen in blue. Sections were differentiated with 1% glacial acetic acid for 2 minutes, dehydrated through graded ethanol and xylene, mounted with xylene-based medium, and stored at room temperature for imaging and analysis. Digital images were captured from all slides using a high-throughput slide-scanner NanoZoomer system (v2.0, Hamamatsu) with a 20× objective and analysed using Nanozoomer digital pathology software. All digital images were processed using the ImageJ/Fiji software. The blue-stained (fibrotic) areas were quantified and analyzed, and the area percentages were determined to reflect variations in the severity of fibrosis. The fibrosis scoring method used in this study is detailed in the previous literature ^75, 76, 77^.

### Statistical Analysis

Differentially expressed genes (DEGs) in bulk RNA-seq and single-cell datasets were identified using *p*-value calculations by the default methods provided by their respective R packages. In both *in vitro* and *in vivo* experiments, statistical differences between different groups were assessed using a two-tailed unpaired Student’s t-test, one-way ANOVA, and two-way ANOVA with significance defined as *p*-value below 0.05. Graphs and statistical analyses were generated using GraphPad Prism 8 (GraphPad Software, www.graphpad.com). Graphical representations were created with BioRender.com.

## Acknowledgements

We would like to thank Louise Turvill, Chief Histologist, Histology facility, Dept. of Pathology, for her assistance with H&E and Masson’s Trichrome staining. We thank Joana Cerveira, Sameen Khan, and Mercedes Cabrera Jarana, Cytometry facility, Dept. of Pathology for their help with flow cytometry. We also thank James Dooley and Adrian Liston for supporting *in vivo* aspects of the study and for their valuable feedback. We thank Bartlomiej Swiatczak for his valuable feedback and comments. We appreciate the support and animal husbandry provided by the UBS animal facility at the Gurdon Institute. Additionally, we acknowledge Novogene (https://www.novogene.com/eu-en/) for their expertise and support in conducting our transcriptomic analyses.

## Author contribution

HAMH: Act as a project lead, conceptualized the study, designed and performed experiments. Analysed, assembled, and visualised data. Wrote the manuscript. SKS: Performed and contributed to the mice experiments, sickness scoring, tissue collection and processing, flow cytometry staining and data acquisition. CV: Contributed to mice experiments, tissue collection & processing, RT-qPCR, and immunofluorescence staining. JP: contributed to mice experiments, tissue collection. SR: scRNA-seq analyses and visualization. FAZA: Histopathological examination, quantification, and scoring analysis. JY: Bulk RNA-seq analysis, visualization, analyses, assembled and visualised data. DH: Supervised JY. EK: mice experiment, sickness scoring, tissue collection, and processing. NM, and YA: LPS administration. QZ: contributed to bioinformatic analysis. BM: Supervised the study. Led the team, reviewed the manuscript, conceptualized the study, fund acquisition, and resource & team management. All authors read and approved the draft manuscript before submission.

## Data and materials availability

Newly generated RNA sequencing data of this study will be submitted to the public repository during the review process. All data will be provided as a Source Data with this paper before the formal acceptance of the paper.

## Additional information

Extended Data Fig. 1

Extended Data Fig. 2

Extended Data Fig. 3

Extended Data Fig. 4

Extended Data Fig. 5

Supplementary Table 1

Supplementary Table 2

Supplementary Table 3

---

**Extended Data Fig. 1.**
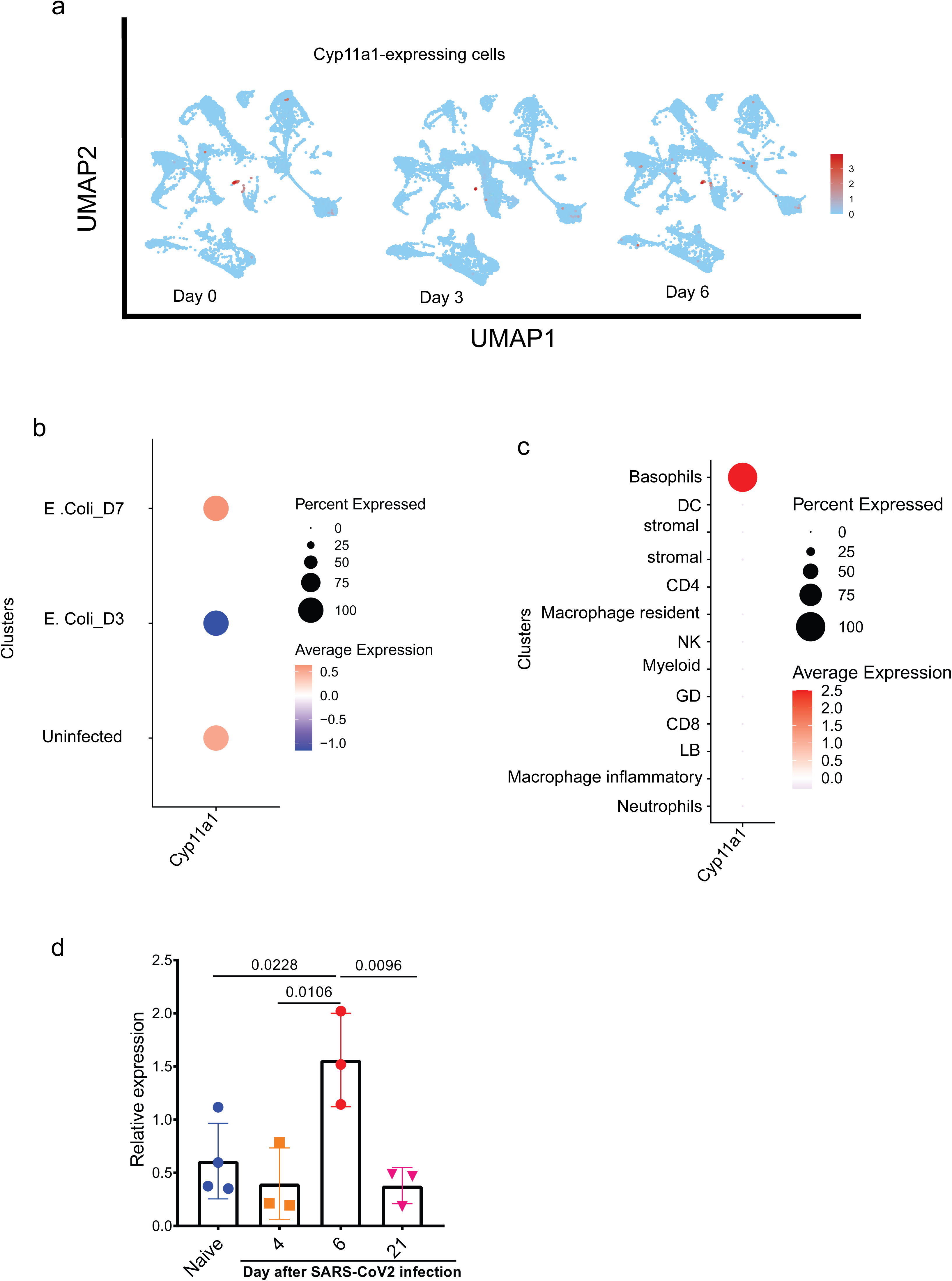
Cyp11a1 expression following respiratory infection. **a,** UMAP projection shows the expression of Cyp11a1 gene across all cells. **b & c,** reanalyzed single-cell transcriptomics data from lung of *E. coli* induced pneumonia model (PRJEB52332). **b**, dot plot shows the proportion and average expression of Cyp11a1 across all immune cells in the lung at day 0, day 3, and day 7 post infection. **c**, dot plot shows the proportion and average expression of Cyp11a1 in individual cells across all time points post infection. **d**, reanalyzed bulk RNA sequencing data generated from SARS-CoV-2–infected mouse lungs show the relative expression of Cyp11a1 in the lung in naive mice and at days 4, 6, and 21 post infection. Bars indicate the mean ± s.d. The *P* value was calculated using a one-way ANOVA.

**Extended Data Fig. 2.**
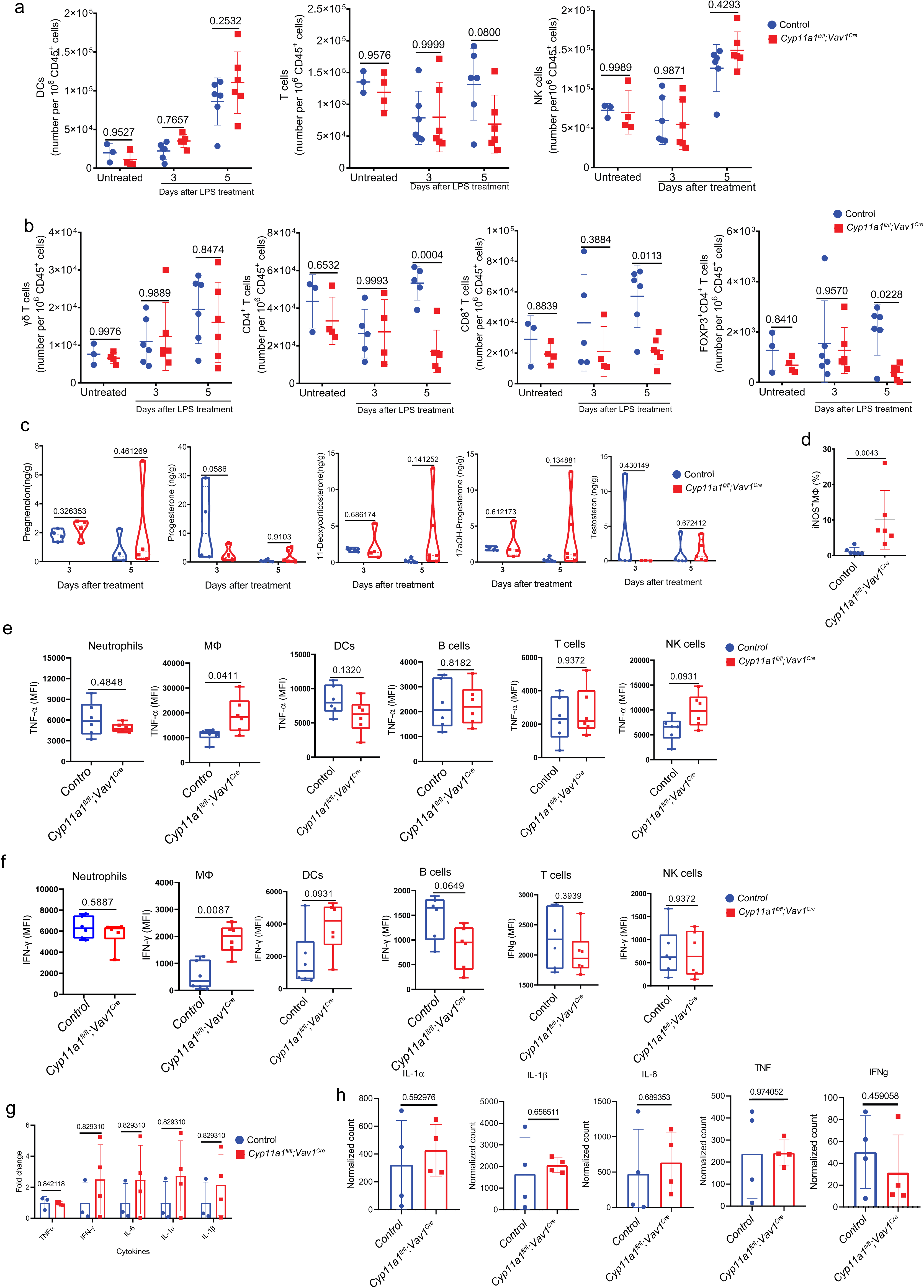
Deletion of Cyp11a1 in immune cells compromises resolution of inflammation following LPS-induce lung injury. **a & b,** flow cytometry counts of DCs, T cells, NK cells (**a**) and T cell subsets (**b**) in the lung at day 0, 3, and 5 post LPS treatment. **c**, profiling different steroids in the lung at day 3 and 5 post LPS treatment. **d**, percentage of iNOS^+^MΦs in the lung at day at day 5 post LPS treatment detected by flow cytometry. **e**, expression of TNF-α in neutrophils, total MΦs, DCs, B cells, T cells, and NK cells at day 5 post LPS treatment detected by flow cytometry. **f**, expression of IFN-γ in neutrophils, total MΦs, DCs, B cells, T cells, and NK cells at day 5 post LPS treatment detected by flow cytometry. **g**, Fold change in the expression of inflammatory cytokines between *Cyp11a1^fl/fl^;Vav1^Cre^* and control mice at steady state detected by RT-qPCR. **h**, normalized count of the expression of inflammatory cytokines between *Cyp11a1^fl/fl^;Vav1^Cre^*and control mice at day 3 post LPS-treatment detected by bulk RNA-sequencing. Bars indicate the mean ± s.d. The *P* value was calculated using a two-way ANOVA (a-c) and two-tailed unpaired *t*-test (d-h).

**Extended Data Fig. 3.**
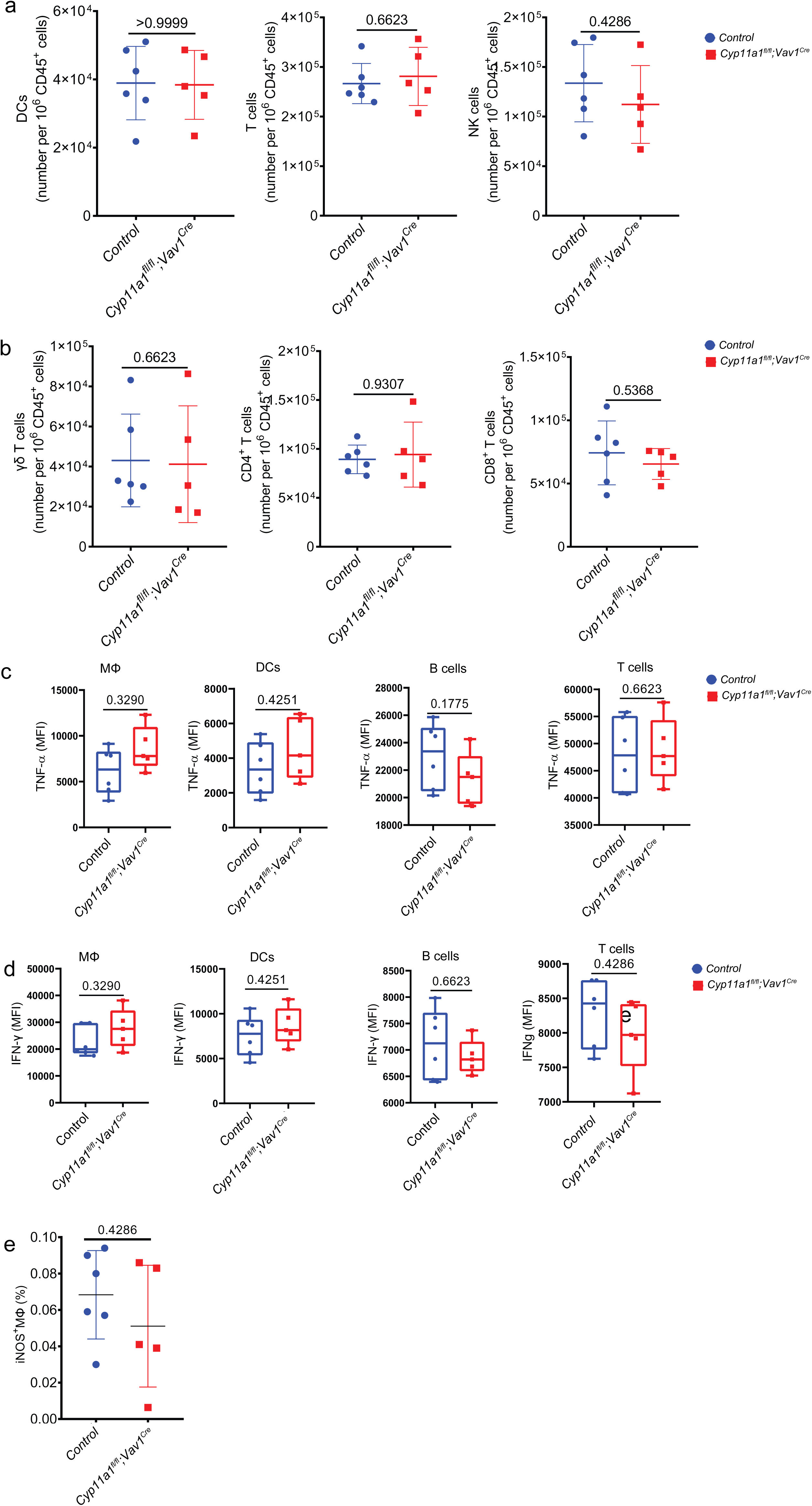
Deletion of Cyp11a1 in immune cells compromises lung repair and promote tissue fibrosis following LPS-induce lung injury. **a & b,** flow cytometry counts of DCs, T cells, NK cells (**a**) and T cell subsets (**b**) in the lung at day 14 post LPS treatment. **c & d**, expression of TNF-α (**c**) and IFN-γ (**d**) in total MΦs, DCs, B cells, and T cells at day 5 post LPS treatment detected by flow cytometry. **e**, percentage of iNOS^+^MΦs in the lung at day at day 14 post LPS treatment detected by flow cytometry. Bars indicate the mean ± s.d. The *P* value was calculated using a two-tailed unpaired *t*-test.

**Extended Data Fig. 4.**
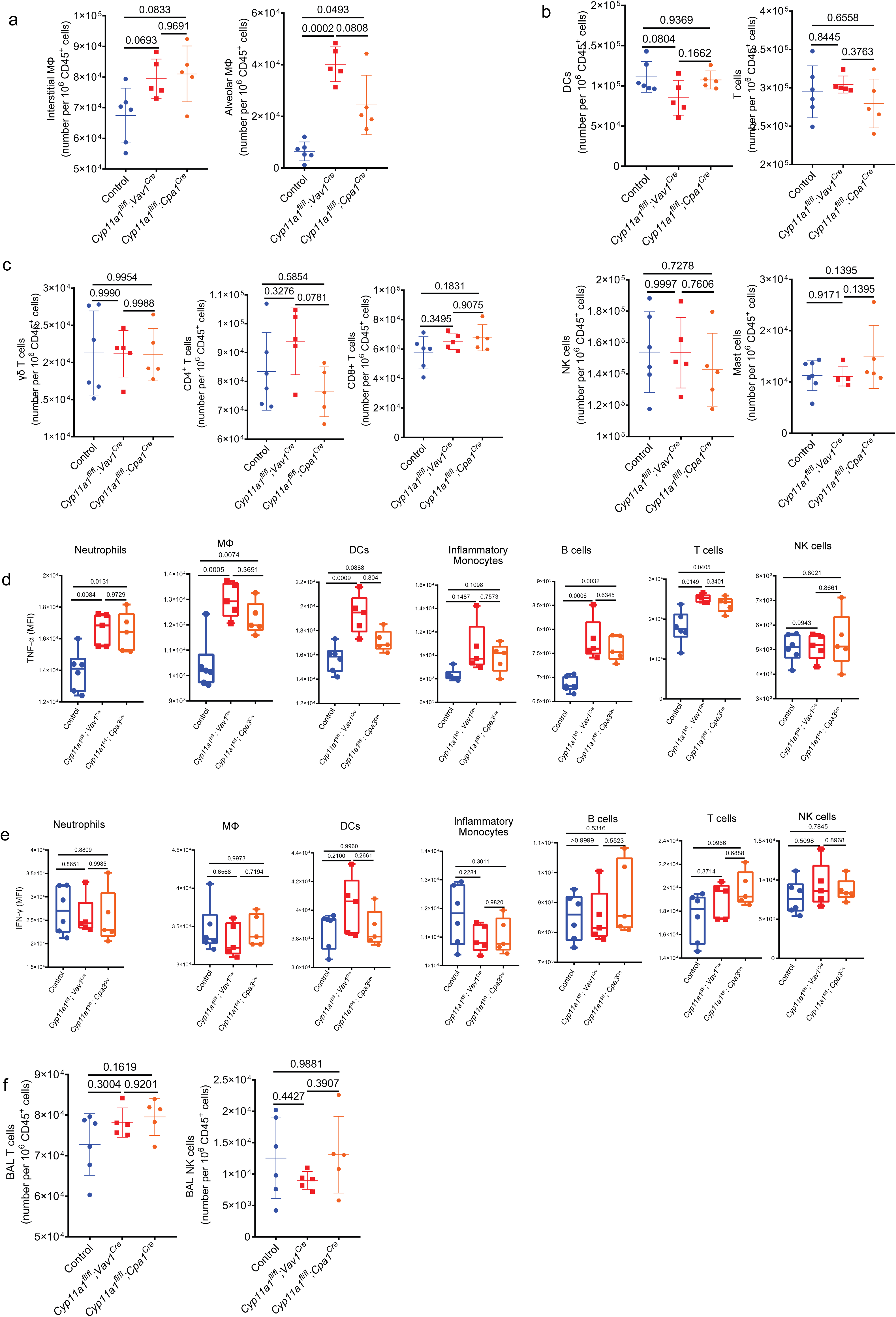
Deletion of Cyp11a1 in basophils compromises resolution of inflammation following LPS-induced lung injury. **a-c,** flow cytometry counts of Ims & Ams (**a**), DCs, T cells, NK cells, and mast cells (**b**) and T cells subsets (**c**) in the lung at day 5 post LPS treatment. **d**,**e,** expression of TNF-α (**d**) and IFN-γ (**e**) in neutrophils, total MΦs, DCs, inflammatory monocytes, B cells, T cells, and NK cells at day 5 post LPS treatment detected by flow cytometry. **f**, flow cytometry counts of T, and NK cells in BAL fluid at day 5 post LPS treatment. Bars indicate the mean ± s.d. The *P* value was calculated using a one-way ANOVA.

**Extended Data Fig. 5.**
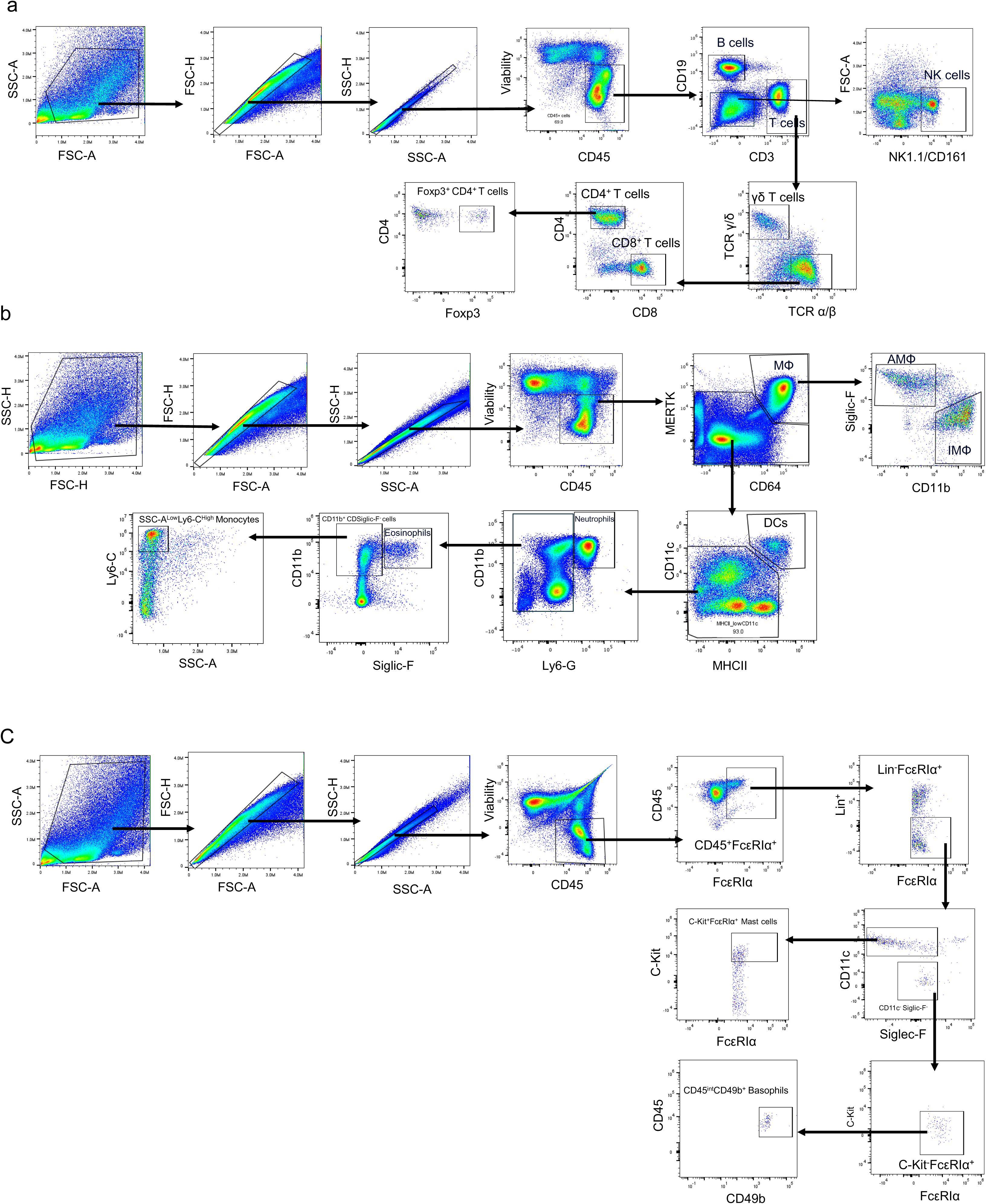
Dot-plots and gating strategies for multi-color flow cytometry. Representative flow cytometry dot-plots and gating strategies for quantification of (**a**) T cell subsets and B cells, (**b**) myeloid cells, and (**c**) basophils and mast cells.

**Supplementary Table 1:** Sickness scoring system.

| **Body weight** |  |
| --- | --- |
| 0 | Normal |
| 1 | 1- 5% |
| 2 | 5-10% |
| 3 | 10-15% |
| 4 | More than 15% |
| **Body posture** |  |
| 0 | Normal body posture |
| 1 | Mild hunch |
| 2 | Moderate hunch |
| 3 | severe hunch |
| **Activity** |  |
| 0 | Normal |
| 1 | Slightly reduced activity |
| 2 | Slow |
| 3 | Very slow |
| 4 | Not moving at all |
| 5 | Dead |
| **Breathing** |  |
| 0 | Normal |
| 2 | Slightly elevated |
| 3 | Distinctly elevated |
| **Coat** |  |
| 0 | Normal |
| 1 | Ruffled coat |
| 2 | Ruffled and not clean |

**Supplementary Table 2:**
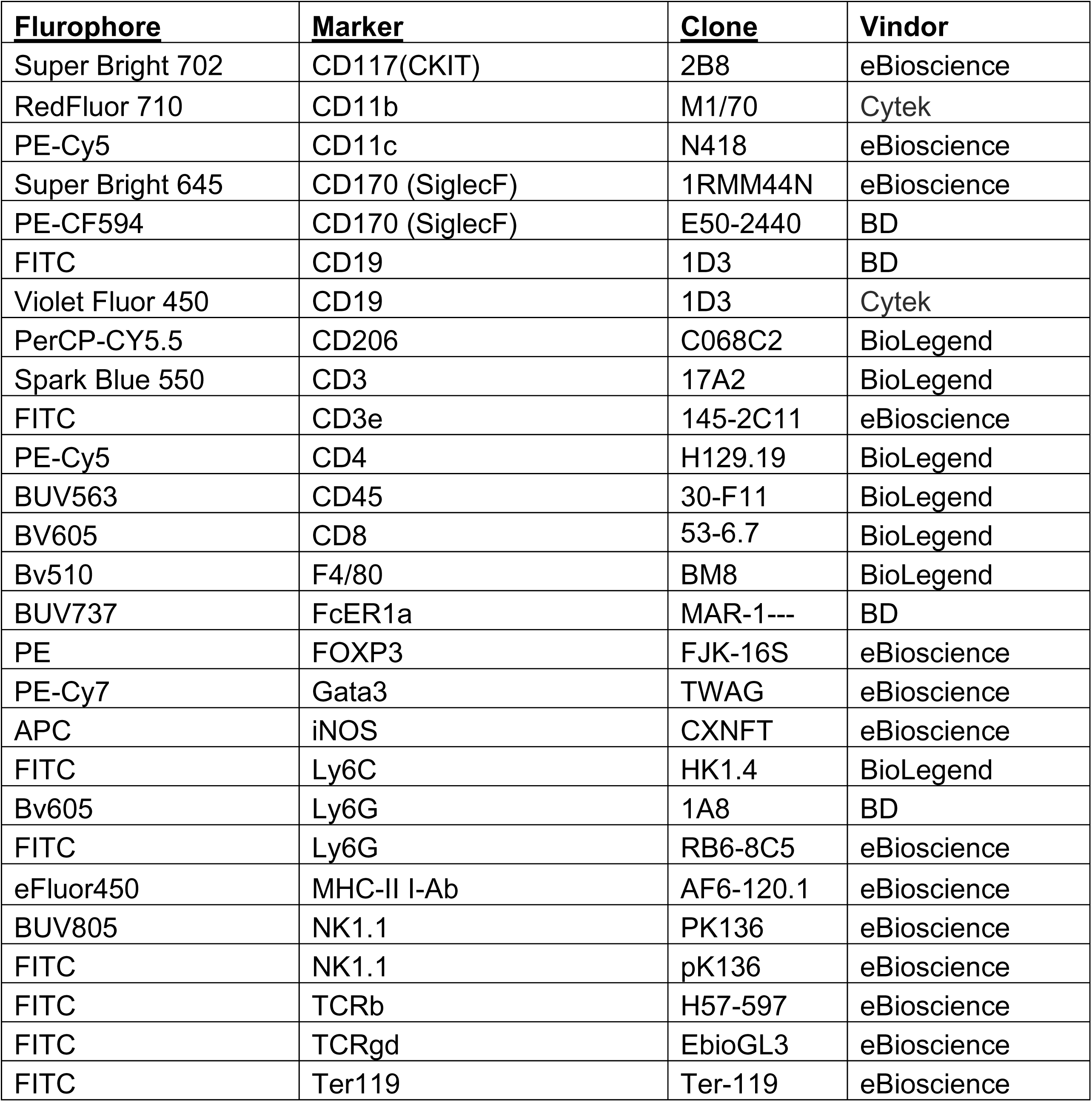
List of antibodies for flow cytometry and Immunofluorescence.

| **Fluorophore** | **Marker** | **Clone** | **Vindor** |
| --- | --- | --- | --- |
| Super Bright 702 | CD117(CKIT) | 2B8 | eBioscience |
| RedFluor 710 | CD11b | M1/70 | Cytek |
| PE-Cy5 | CD11c | N418 | eBioscience |
| Super Bright 645 | CD170 (SiglecF) | 1RMM44N | eBioscience |
| PE-CF594 | CD170 (SiglecF) | E50-2440 | BD |
| FITC | CD19 | 1D3 | BD |
| Violet Fluor 450 | CD19 | 1D3 | Cytek |
| PerCP-CY5.5 | CD206 | C068C2 | BioLegend |
| Spark Blue 550 | CD3 | 17A2 | BioLegend |
| FITC | CD3e | 145-2C11 | eBioscience |
| PE-Cy5 | CD4 | H129.19 | BioLegend |
| BUV563 | CD45 | 30-F11 | BioLegend |
| BV605 | CD8 | 53-6.7 | BioLegend |
| Bv510 | F4/80 | BM8 | BioLegend |
| BUV737 | FcER1a | MAR-1--- | BD |
| PE | FOXP3 | FJK-16S | eBioscience |
| PE-Cy7 | Gata3 | TWAG | eBioscience |
| APC | iNOS | CXNFT | eBioscience |
| FITC | Ly6C | HK1.4 | BioLegend |
| Bv605 | Ly6G | 1A8 | BD |
| FITC | Ly6G | RB6-8C5 | eBioscience |
| eFluor450 | MHC-II I-Ab | AF6-120.1 | eBioscience |
| BUV805 | NK1.1 | PK136 | eBioscience |
| FITC | NK1.1 | pK136 | eBioscience |
| FITC | TCRb | H57-597 | eBioscience |
| FITC | TCRgd | EbioGL3 | eBioscience |
| FITC | Ter119 | Ter-119 | eBioscience |

**Supplementary Table 3:**
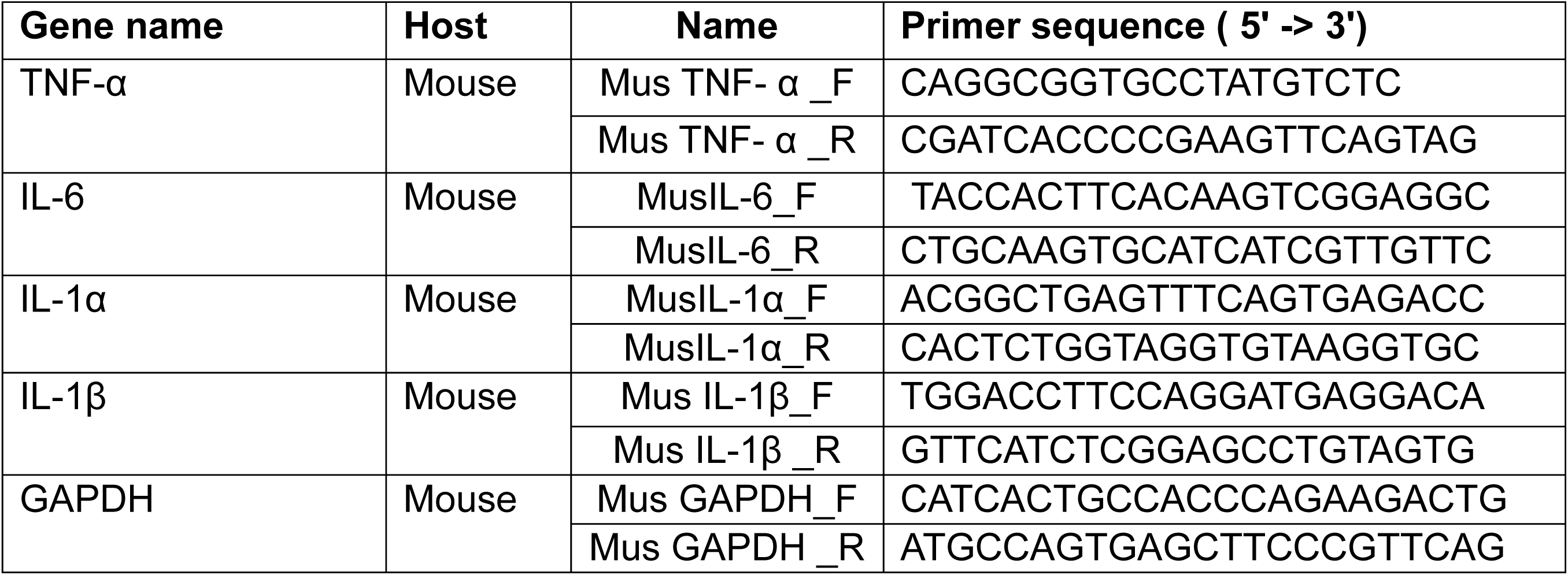
RT-qPCR Primer list.

